# Larval density in the invasive *Drosophila suzukii*: immediate and delayed effects on life-history traits

**DOI:** 10.1101/2022.10.06.511120

**Authors:** Alicia Reyes-Ramírez, Zaïnab Belgaidi, Patricia Gibert, Thomas Pommier, Aurélie Siberchicot, Laurence Mouton, Emmanuel Desouhant

## Abstract

The immediate and delayed effects of density are key in determining population dynamics, since they can positively or negatively affect the fitness of individuals. These effects have great relevance for polyphagous insects for which immature stages develop within a single site of finite feeding resources. *Drosophila suzukii* is a crop pest that induces severe economic losses for agricultural production, however little is known about the effects of density on its life-history traits. In the present study, we (i) investigated the egg distribution resulting from females’ egg-laying strategy and (ii) tested the immediate and delayed effects of larval density on emergence rate, development time, sex ratio of offspring, fecundity and adult size (a range of 1 to 50 larvae was used). We showed that most of fruits contain several eggs and aggregate of eggs of high density can be found in some fruits. This high density has no immediate effects on the emergence rate, but has effect on larval developmental time. This trait was involved in a trade-off with adult life-history traits: the larval development was reduced as larval density increased, but smaller and less fertile adults were produced. Our results should help to better understand the population dynamics of this species and to develop more successful control programs.

## Introduction

Density-dependence is a key driver of demographic parameters (Hixon & Johnson, 2009). It may have positive or negative effects on a population growth rate (Mueller, 1997). These effects can participate in the early stages of the invasion process by establishing minimum population sizes that, when exceeded, promote the spread of pests (Fagan et al., 2002). Among these invasive, pest or biocontrol auxiliaries species, many are insects (Ferguson & Joly, 2002; Pickens, 2007). Density-dependent effects are therefore common features that define invasive or pest population dynamics’ as well as the success of biocontrol programs.

Density-dependent effects on demographic parameters emerge from the combination of effects occurring at lower levels of biological organization (Mueller, 1997; Ponton & Morimoto, 2020). Effects observed at the individual level can be classified into immediate and delayed effects. Immediate effects typically occur at immature stages and are associated with resource acquisition (Dethier, 1959; Putman, 1977). For many insect species, nutritional resources may be in limited quantity when females lay their eggs in a finite volume of food such as fruits or seeds, and when the whole immature development occurs in a given seed/fruit. Therefore, female oviposition strategies affect early developmental conditions and thus larval fate as well as adult traits (Nestel et al., 2016). Delayed effects are the result of trade-offs that emerge in response to density-dependence conditions during development and that are expressed at later life stages (Agnew et al., 2002). Both immediate and delayed effects can positively or negatively affect individual fitness (Peters, 2003) by altering life-history traits and promoting trade-offs (Parker & Gilbert, 2018). For example, in the lepidoptera *Sesamia nonagriodes*, the high density experienced during the larval stages does not affect the mortality of juveniles, but extends larval development, and results in reduced pupal weight (Fantinou et al., 2008). In this species, adult fitness is also affected with reduced fecundity and longevity of females reared in high larval densities. In general, individuals that develop in low density environments are predicted to have an advantage in later life due to expecting low levels of competition for resources (the “silver spoon” effect; Angell et al., 2020; Grafen, 1988). For example, in the speckled wood butterfly, *Pararge aegeria*, larvae reared at low densities have higher survival rate, shorter development times and result in bigger adults compared to larvae reared at high densities (Gibbs et al., 2004). Nevertheless, high density may also be beneficial, for example when individuals express a greater performance in defense against predators (Aukema & Raffa, 2004), in cooperative feeding (Denno and Benrey 1997), or when beneficial horizontally transmitting microbiota are present (Correa et al., 2018).

These examples highlight the need to study both the immediate and delayed effects of density to understand and predict population dynamics, especially for pest species, in order to more efficiently manage and control their populations (Alkema et al., 2019). In this study, we focus on *Drosophila suzukii*, a pest of many berry and stone fruit crops in Asia, Europe and America (Dos Santos et al., 2017; Lee et al., 2011). Females possess a serrated ovipositor (Atallah et al., 2014) that allow them to lay eggs in healthy fruits without any wounds, unlike most other Drosophilidae that oviposite on ripe or damaged fruits (Mitsui et al., 2006). This polyphagous fly thus induces severe economic losses for agricultural production (Knapp et al., 2021). Despite its main agricultural impacts, the immediate and delayed effects of density on life-history traits have not been deeply investigated in this species. Few publications indicate that at high larval densities, the weight (Kienzle et al., 2020) and survival (Wang et al., 2019) of adults decrease. A high density can also alter the chemical composition and microbial diversity of the food medium in which larvae developed due to foraging and excretion of conspecifics (Henry et al., 2020).

In the present study, we aimed at characterizing the immediate and delayed effects of larval density on major life history traits of *D. suzukii*. To fulfil this objective, we first investigated how females distribute their eggs in fruits in order to establish a relevant range of larval density per fruit to test the effect of larval densities on larval and imaginal life-history traits. Densities ranging from 1 to 50 larvae were thus tested in two resource volumes mimicking two sizes of fruit. Due to the absence of relevant literature, we had two opposite predictions: high densities have positive effects due to feeding facilitation or negative effects on life-history traits due to larval competition. Likewise, we expected that, according to the larval density and the volume of food resource, developmental time, adult emergence rate, sex ratio of offspring, fecundity and adult size, should change. Finally, we also tested whether the microbial diversity and colony counts changed with the larval density.

## Materials and methods

### Drosophila suzukii line and rearing conditions

We used a *Wolbachia-free* line of *D. suzukii* originated from the Agricultural Entomology Unit of the Edmund Mach Foundation in San Michele All’Adige, Trento Province, Italy (Nikolouli et al., 2020). Before and after the experiments, the absence of *Wolbachia* was checked by PCR (see TableS1 for protocol). The flies were reared on a cornmeal diet containing: 0.9% agar, 5% sugar, 3.3% cornmeal, 1.7% dried yeast, 0.4% nipagine, and maintained in an incubator at constant temperature (22.5 °C) and humidity (60%) with a 12-hours light/dark cycle.

### Oviposition assays

Egg-laying behaviour was observed on blueberries. In a plexiglass box (23.8 x 17.8 x 2 cm), 3 groups of two blueberries (from organic farm) were placed using double-sided tape at equal distance from each other (FigureS1). A piece of sugar agar medium was placed in the center of the box to ensure the nutrition of the flies and hydration. In each box, one 7 days-old mated female was placed for 18h (32 replicates were done) and then the number of eggs per fruit was counted under a binocular loupe.

### Experimental protocol for immediate and delayed effects of larvae density on life-history traits

Effect of larval density was tested using 1, 5, 10, 20 and 50 larvae and two volumes of medium, 2 or 5mL, in 25mL tubes (Eppendorf® Conical Tubes). For each of these 10 combinations of modalities, at least 8 replicates were performed (Figure1). The larval development conditions were standardized by allowing mated females at least one week old to oviposit for 24h. After egg hatching, larvae of the first stage (L1) were collected and then randomly assigned to one of the ten experimental modalities. The experiments were conducted in two temporal blocks.

**Figure1.**
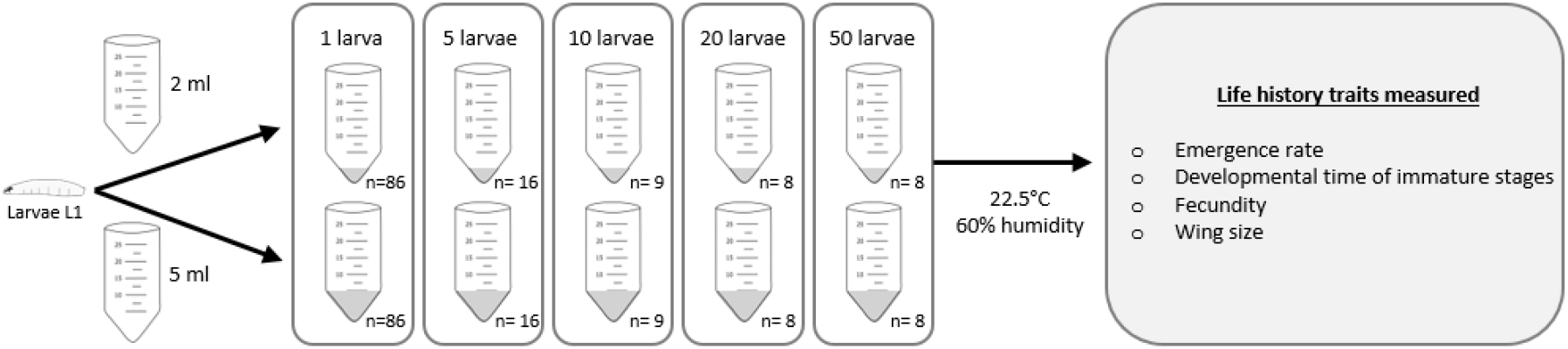
Experimental protocol for testing immediate and delayed effects of larval density on life-history traits.

Several larval and adult life history traits were measured:

### Preimaginal developmental time and adult emergence rate

Tubes were checked twice a day in order to detect adult emergencies. For each tube, the emergence rate was calculated by comparing the number of adult flies with the number of larvae sown. Developmental time was estimated for each individual between L1 and adult stages.

### Potential fecundity

Potential fecundity was assessed on 3-days-old emerging females by dissection of their abdomen in PBS. To put them sleep, females were first placed for at least 30 min in the freezer. The number of mature eggs was counted in the two ovaries (20 females per modality) with a binocular loupe as described in Plantamp et al. (2017).

### Wing length and width

Prior to dissection, the right wing of females, a classical proxy of the adults’ size in *Drosophila* (David et al., 1994), was taken and placed on a microscope slide. Coverslips were sealed using nail polish. Images of the wings were acquired using the AxioVisio 4.8 software on a Zeiss Imager.Z1. microscope. Two measures were performed (see FigureS2) on 20 females per modality: the length corresponds to the distance between the tip of the wing, and the R_4+5_ vein, and width to the distance between the R_2+3_ and the Cu_A1_ veins (https://commons.wikimedia.org/wiki/File:Drosophilidae_wing_veins-1.svg).

### Diversity and quantity of microorganisms

In order to test whether the larval density (and thus feeding and excretion) changes the diversity of bacteria in the food medium, we inoculated medium from the vials where the larvae developed in. Using an inoculation loop, around 10μL of food medium were sampled under sterile conditions and diluted in 100μL of ultrapure water. These mixtures were then streaked on two solid growth media, LB (Lysogeny Broth amended with 5% Agar) and TSA (Tryptone Soy Agar), in 90 mm Petri dishes (8 boxes per modality and per medium, *i.e*. 160 Petri dishes in total). After sealing with parafilm to limit drought and cross-contaminations, the Petri dishes were incubated aerobically for 7 days at 37 °C in the dark. The counts were realized on ¼ randomly chosen part of the Petri dishes at the end of the incubation. Control Petri dishes were incubated to evaluate potential contamination during incubation. All controls remained blank until the end of the experiment.

### Statistical analysis

To test whether, after 18h of oviposition period, the distribution of eggs laid by a female was aggregated or random, we fitted different theoretical distributions (GLMs with Poisson, negative binomial (NB) distributions, and zero-inflated negative binomial (ZINB) distributions, with log and logit link, respectively) to the number of eggs deposited per fruit, and to the number of fruits infested per female. NB and ZINB are usually used to fit aggregated distributions. ZINB is used for overdispersed count variables, allowing to model data with excessive zeros. The AIC values were used to compare the different fitted models.

We tested whether there was a differential oviposition rate between females with two mixed generalized linear models (GLMM) adjusted with a Poisson distribution. For each GLMM, the total number of eggs and the number of infected blueberries were the dependent variables, respectively, while the box was the independent variable. In both models, the date was included as a random factor.

The effects of larval density on emergence rate (*i.e*. the total number of emerging adults per tube) were analyzed with a GLMM (binomial distribution, logit link). We included the volume, the density and the sex of the individuals as independent variables, as well as all the double interaction, and the block as a random factor.

The effects of larval density on the number of microbial colonies per medium plate were analyzed by the means of a GLMM (Poisson distribution, log link). We also included the volume and the growth medium as independent variables, as well as all the double interactions; the tube where the aliquot was taken was included as a random factor. A Tukey test was used for comparisons between treatments.

To test the effect of density and volume on life-history traits, four different GLMs were performed. Because of the unbalanced designs, we performed type-III analysis of variance (Shaw & Mitchell-Olds, 1993) for each of these models. Development time, fecundity and size of the wings (length and width) were used as dependent variables, while volume, density and block corresponded to the independent variables. In the case of development time, the sex of the emerging individuals was also used as an independent variable. We compared treatments using the post-hoc Least Significant Difference (LSD) test with Benjamini-Hochberg (1995) procedure, whereby a separate analysis for each treatment and corresponding interactions are obtained (Engqvist, 2005). In the case of density, the control was 1 larva. Pairwise Spearman correlation was calculated between wings’ length and wings’ width. All analyses were performed in R version 4.0.2 (Team, 2020) with the “emmeans” (Lenth & Lenth, 2018), “multcompView” (Graves et al., 2015) and “car” (Fox & Weisberg, 2019) packages.

## Results

### Oviposition assays

The objective of this first experiment was to have an estimation of the range of larval density per fruit. Globally, 44.27% of the blueberries were infested (per box 2.65±1.75). The number of eggs per infested fruit varied from 1 to 11 (1.34±0.4). 31.77% of females oviposited more than one egg per fruit.

ZINB turned out to be the best model to explain the distribution of the number of eggs per fruit (TableS2), showing that the distribution of eggs was in aggregates with an extra number of non-infested fruits than expected under classical negative binomial distribution (z=6.47, p<0.001; FigureS3). No significant differences (χ^2^_1_=2.887, p=0.089) between females were found in the number of eggs deposited per box (1 to 24 eggs, 8.03±6.34). We draw the same conclusion for the number of blueberries infected per box (χ^2^_1_=2.155, p=0.142).

### Immediate effects of larval density

#### Effect of larval density and resource volume on adult emergence

The average emergence rate was 0.59±0.06. Neither larval density nor the volume of food affected emergence rate (χ^2^_1_=0.277, p=0.598; χ^2^_1_=2.444, p=0.118 respectively; Figure2A and FigureS4A), and there was no significant interaction between these two variables (χ^2^_1_=3.198, p=0.073). Fly emergence did not vary with the sex of the emergent individuals (χ^2^_1_=0.167, p=0.682), and we did not detect interactions between sex and density (χ^2^_1_=1.992, p=0.158) or between sex and volume (χ^2^_1_=0.441, p=0.506).

**Figure2.**
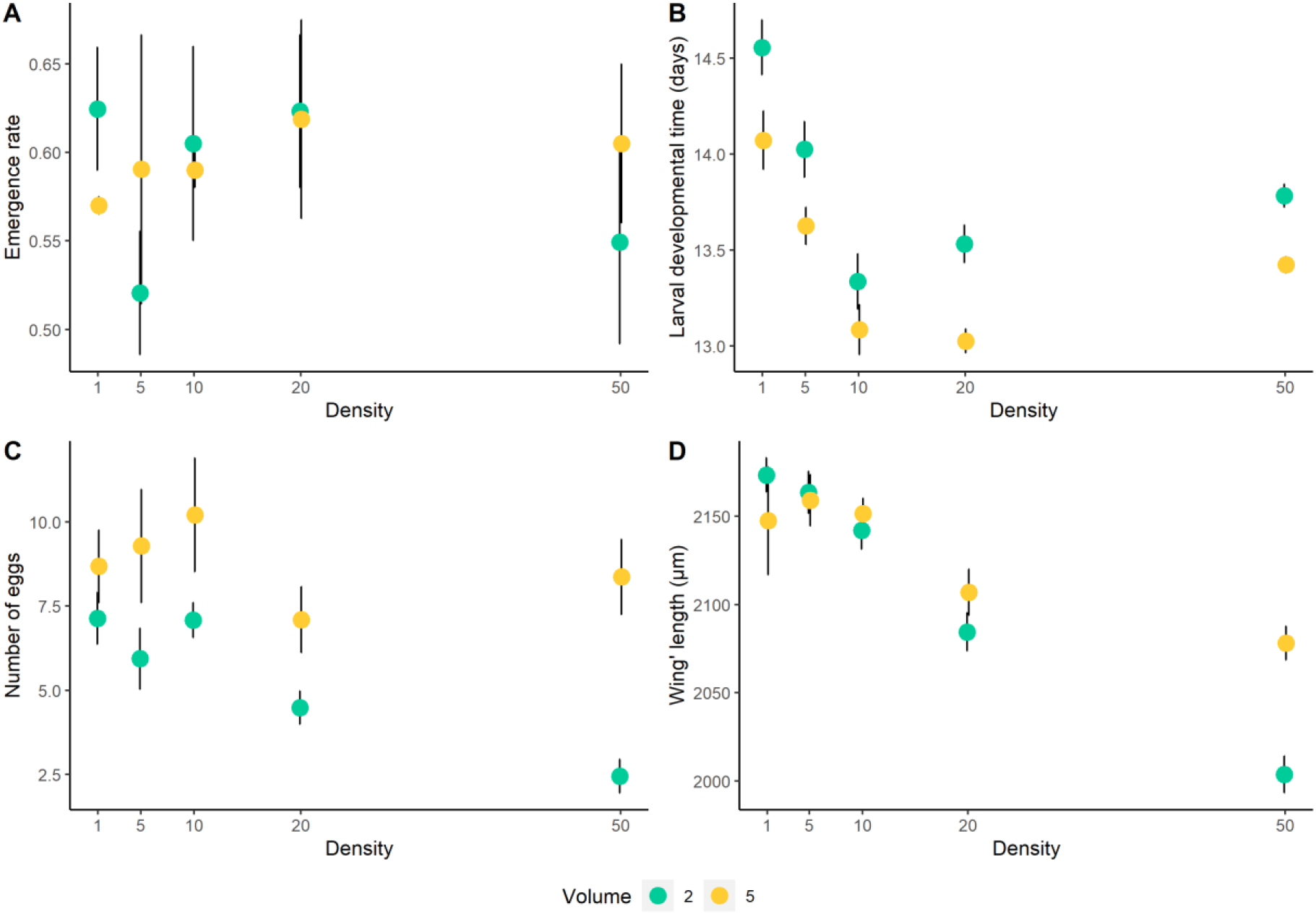
Effect of larval density and resource volume (2 (green) and 5 (yellow) mL of food medium) on immediate (A, B) and delayed (C, D) life-history traits (mean ± SE) of *D. suzukii*. Panel (A) shows emergence rate. Panel (B) shows the developmental time for larvae to emerge in days. Panel (C) shows fecundity measured as the number of eggs. Panel (D) shows wings’ length.

#### Effect of larval density and resource volume on larval development time

The larval development time was affected by volume of resource available for larval feeding (F_1,940_=50.632, p<0.001), density (F_4,940_=32.437, p<0.001), sex (F_1,940_=47.53, p<0.001) and block (F_1,940_=82.843, p<0.001; TableS3). In general, individuals that have grown in the lowest larval density (*i.e*. 1 larva) took more days (0.83±0.21) to develop compared to the other densities (Figure2B). Also, the emerging females took more days to develop than males (mean difference=0.45±0.03; FigureS6). The interaction between volume and block (F_1,940_=37.803, p<0.001; TableS4) and density and block (F_4,940_=10.778, p<0.001; TableS5) were significant. The individuals raised in the block1 took longer time to develop than individuals tested in block2 in both resource volume (FigureS5B).

The same pattern was observed in the interaction with density, excepted for density 50 where developmental times were the same for the two blocks (TableS6, FigureS4B).

#### Effect of larval density and resource volume on microbial diversity and colony counts

Whatever the larval density and the volume of resources, all colonies harbored an identical morphology, suggesting a limited diversity in all modalities (FigureS7).

However, the colony counts differed with a significant interaction between larval density and the bacterial growth medium (χ^2^_4_=112.858, p<0.001; TableS7, FigureS8). The number of colonies (174.23±140.07) increased with the density of larvae (TableS8, Figure3), and globally colonies were more numerous in the LB medium as compared to the TSA medium (TableS8, S9). Likewise, the interaction between the volume of food and the growth medium was significant (χ^2^_1_=21.323, p<0.001), mainly due to a fewer number of colonies grown on TSA medium compared to LB medium in samples from 2 mL of resources.

**Figure3.**
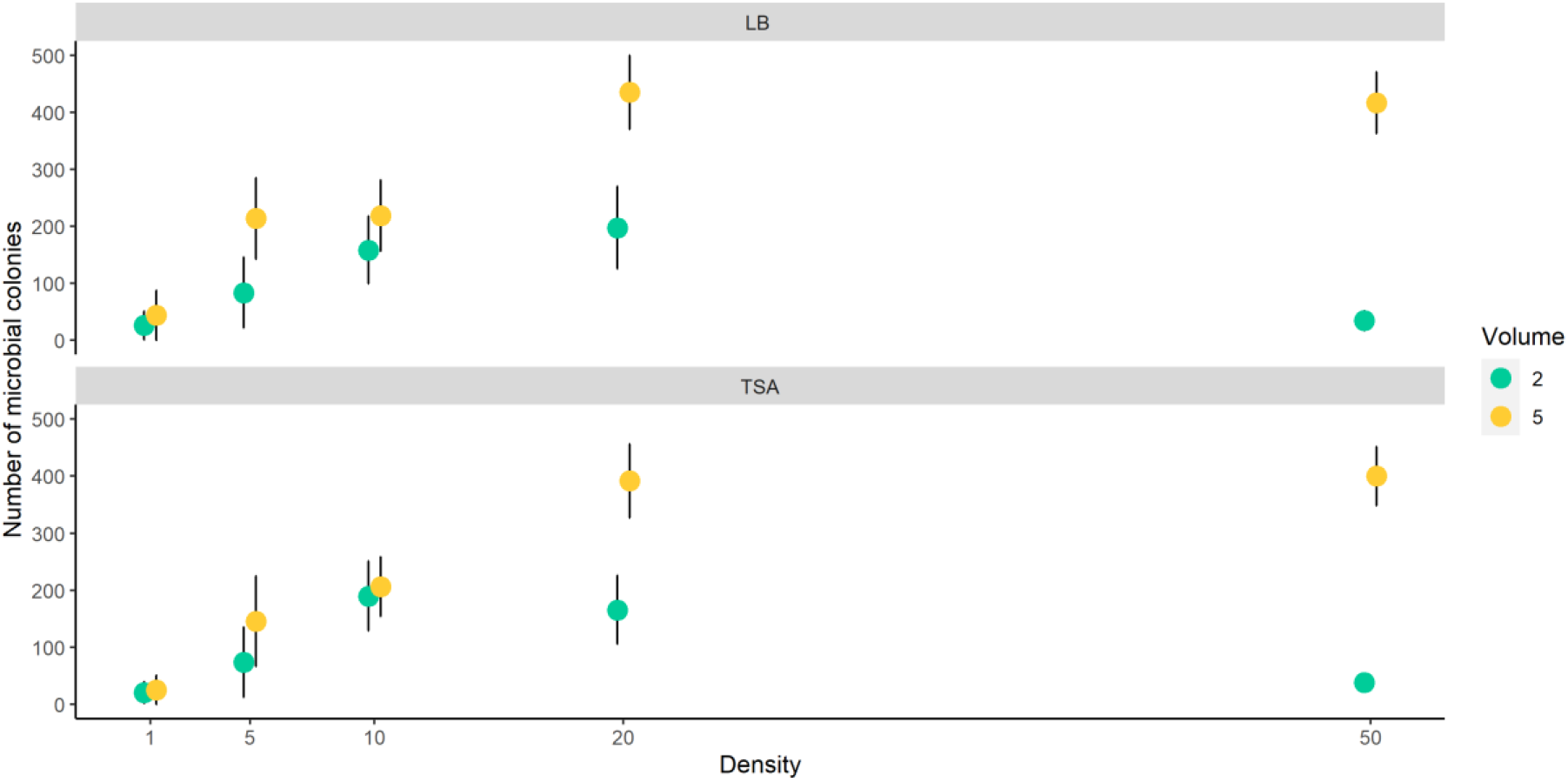
Effect of larval density and resource volume (2 (green) and 5 (yellow) mL of food) on the number of microbial colonies (mean ± SE) present in the medium where *D. suzukii* larvae develop. Above are the colonies grown in LB medium and below those grown in TSA medium.

### Delayed effects of larval density

#### Effect of larval density and resource volume on female fecundity

Females developed in low density laid more eggs (7.91±4.62) compared to those raised at high densities (5.41±4.31). Both, larval density (F_4,241_=4.246, p<0.01; Figure2C) and volume of resources (F_1,241_=27.493, p<0.001) have an impact on fecundity. However, there were differences between blocks (F_1,241_=4.525, p<0.05; TableS10; FigureS4C) with a significant interaction (F_4,241_=2.868, p<0.05; TableS11): at density 5, the individuals that developed in block 2 had more eggs than those that developed in block 1 (z=-3.339, p<0.001; TableS12, FigureS4C).

#### Effect of larval density and resource volume on wing length and width

As the length and the width of the wings were positively correlated (Spearman’s r255=0.84, p<0.001, FigureS9), we presented only results of the length (FigureS10 and see sup mat for width TableS17-TS20). Globally, individuals that have developed with few larvae have a wider wing length than the other ones (F_4,241_=31.408, p<0.001). However, there is a significant interaction between larval density and resource volume (F_4,241_=4.756, p<0.01; TableS13, TableS14, Figure2D) which is mainly due to the differences at density 50: the individuals that had grown in 2 mL emerged with the smallest wings which is the contrary at the other larvae densities (z=-4.17, p<0.0001; TableS15). The interaction between volume and block was also significant (F_1,241_=26.586, p<0.001; TableS16, FigureS5D), but there were no differences between the blocks (F_4,241_=0.256, p=0.61).

## Discussion

Our study gives insights into the range of eggs or larvae that can be found in fruits infested by *D. suzukii*, and the potential effects of the larval density on major life-history traits of this pest and their related trade-offs.

In phytophagous species, for which immature develop within a finite volume of resources (e.g. fruits or seeds), mothers’ oviposition strategy determines the fate of offspring and their fitness (Doak et al., 2006). Our results showed that the oviposition strategy of *D. suzukii* females results in an aggregative distribution of eggs in fruits: until 11 eggs have been laid by one female in the same blueberry, with an average of more than 2.5 eggs per infested fruit. This suggests that, in conditions where fruits could be limiting (for instance in the beginning of the fruit season or in greenhouses cultures), high densities of immature in a given fruit is likely. It is surprising, that for this major pest species, the density of larvae per fruit in the field is still unknown. Only indirect measures are available (Elsensohn et al., 2021) and confirm the possibility of high larval density per fruit (e.g. per berry 4.2±1.3 in raspberries (Burrack et al., 2013) and 2.6±0.8 in mulberries (Yu et al., 2013)). However, our results are not informative about the oviposition strategy *sensu stricto* (*i.e*. choice of fruits, number of eggs laid at each oviposition bout, sequence of eggs deposition). Indeed, as shown by Desouhant et al. (1998) (on the chestnut weevils, *Curculio elephas*), different oviposition strategies (random, aggregate or uniform) can lead to aggregated distributions of eggs. However, our results strongly suggest that, at least in our lab conditions (*i.e*. few available fruits), a *D. suzukii* female does not avoid fruits already containing its own eggs.

Usually larval density affects immediately lifespan and other fitness traits such as development time like in *D. melanogaster* (Horváth & Kalinka, 2016). Here, we observed contrasted effects of density on larval life history traits. We did not find any change in preimaginal survival (from L1 to adult emergence) between densities or between sexes. This conclusion is valid regardless of the resource volume (mimicking two sizes of fruits) while we expected an increase of negative density dependent effects in the small resource volume. In contrast, the larval development time of flies was negatively affected when density increased: *D. suzukii* larvae raised in high densities developed faster. Two mutually non-exclusive hypotheses could explain this result. First, a shorter developmental time could result from facilitation effect. The food medium is more intensively digested, allowing an increase in food intake rate that results in a reduction of the developmental time. Feeding facilitation and short development has been documented in the Queensland fruit fly *Bactrocera tryoni* (Morimoto et al., 2018). Second, a faster development could allow escaping competition and avoid mortality driven by the risk of running out of resources before metamorphosis, as shown in the dung fly *Scathophaga stercoraria* (Blanckenhorn, 1999). However, in our experiments, larvae reared in the smallest volume of food (2 mL, *i.e*. the highest competition intensity) took more days to reach adult stage than those reared in 5 mL, regardless of larval density. This clearly means that, when facing to a greater intra-specific competition intensity (due to a reduction of available resource), the larvae strategy is to compensate for reduced nutrient intake by increasing their development time. Several species of phytophagous insects display this strategy with an extension of the larval period, potentially allowing through an extension of the feeding time to reach a size threshold compatible with metamorphosis and viable physiological conditions (Yang et al., 2015). This developmental plasticity (Mackay, 2001) is expected to be selected for when immature have no opportunity to find food resource outside the oviposition site chosen by their mother, such as in *D. suzukii*. A similar pattern was observed in the tropical butterfly *Bicyclus anynana*. Its larval development time was also reduced when the larvae were reared at high densities, but the development time was prolonged when experiencing food stress, suggesting more time available for feeding (Bauerfeind & Fischer, 2005). At last, we showed that males emerged before females, but the sex ratio was not affected. The sex-specific differences in development time, where females take more days to reach the adult stage, may be associated with maximizing the number of matings for males (protandry) (Teder et al., 2021). In addition, a longer timing of maturation would allow females to reach a bigger size, which entails an advantage in their fecundity (Honěk, 1993; Teder et al., 2021).

Larval density also had an immediate effect on the bacterial community of the larval food medium: though apparent diversity did not change, the number of bacterial colonies increased with the larval density, without any negative effect on the preimaginal survival. Moreover, as we did not observe any changes in bacterial morphotypes according to the different larval densities tested, a confounding effect due to the appearance of another bacterial type is not to be considered. This indicates that the changes in life traits observed are indeed due to the density. A complete picture of the potential interactions between density effects and the emergence of different bacterial or fungal types, as would occur in natural conditions, would require complementary experiments and more discriminating approaches (e.g. using molecular affiliations). They would also permit to test the susceptibility of insects to bacteria in their environment.

Density effects detected on immature stages had also consequences on the adult life-history traits. One of the most remarkable effects of crowding during larval development is the decline of female fecundity. In our study, the females that experienced high larval densities had a lower number of mature eggs present in their ovarioles. This negative impact is observed in numerous species (see Vamosi & Lesack, 2007). As observed in other species (e.g. in *C. vomitoria*, (Ireland & Turner, 2006); and in *B. tryoni*, (Morimoto et al., 2019)), in *D. suzukii*, the decrease in the number of eggs produced is positively correlated to a reduction of the wing size, a proxy of body size (Sokoloff, 1966). Our study also corroborates that body size is an important determinant of female fecundity (see also Honěk, 1993; Leather, 2018).

The results presented above showed a phenotypic trade-off between larval and adult life-history traits. High larval density led immature to develop faster as the expense of adults’ size and fecundity.

Throughout the genus *Drosophila*, this trade-off between juvenile developmental rate and adult viability is described (Prasad et al., 2000). For example, *D. melanogaster* has an antagonistic pleiotropy between developmental rate and early- and late-life survival (Chippindale et al., 2004). We assume that, at high density, female fitness is negatively affected by a reduction in flight ability and thus in the search for suitable oviposition sites, as well as by the risk of becoming egg-limited due to reduced egg load (Rosenheim, 2011). This reasoning implies a negative correlation between size and reproductive success in the field that is not always proven (Ellers et al., 1998; West et al., 1996). In addition, an estimate of the impact of larval density on adult longevity would be relevant to an accurate estimation of density dependent effects on fitness.

## Conclusion

The females of *D. suzukii* laid their eggs in an aggregate distribution which promotes crowding of the larvae. Contrary to expectations, preimaginal survival to adult emergence was not affected. However, larval development time shortened as density increased and resources became more limited. Furthermore, rearing at high larval densities negatively affected the fitness of adults which were smaller and had reduced fecundity. Our results support the existence of trade-off between larvae and adult life-history traits. A greater understanding of the effect of density on the population dynamics of this pest may result in more successful alternative control measures.

## Acknowledgments

This work was supported by the ANR (ANR-19-CE32-0010), which also awarded A. Reyes-Ramírez post-doctoral fellowship. We also thank X. Fauvergue, N. Ris, A. Auguste and L. van Oudenhove for useful discussion on data.

