## Supplementary material for "Larval density in the invasive *Drosophila suzukii*: immediate and delayed effects on life-history traits"

Table S1. PCR primers and conditions used to check for the presence of *Wolbachia*

| Primers | Primer sequences | Annealing temperature/ Product size | References |
|---|---|---|---|
| **81F**<br>**2R** | 5' – TGGTCCAATAAGTGATGAAGAAAC – 3'<br>5'- CAGCAATTTCAGGATTAG -3' | 54°C / 290bp | Braig et al. 1998<br>Henri & Mouton 2012 |

DNA from single individuals was extracted using the NucleoSpin Tissue kit (Macherey-Nagel). *Wolbachia* detection was performed using *Wolbachia* specific primers that amplify a part of the *Wolbachia surface protein* gene. PCR reactions were performed in 10µL volumes containing 500nM primers, 1X Precision Melt Supermix (Biorad®) containing dNTPs, iTaq™ DNA polymerase, MgCl2, EvaGreen dye, stabilizers) and 2µL of DNA diluted to one tenth. Cycling conditions were 95°C for 2 min, then 30sec at 95°C, 30sec at 54°C and 30 sec at 72°C for 40 cycles, followed by 30 sec at 95°C and 1min at 60°C (Biorad CFX96).

TableS2. AIC values for the different models used in the distribution of oviposited eggs in blueberries. Comparisons were made between ordinary least squares (OLS), Poisson, negative binomial and the zero-inflated negative binomial regression (ZINB).

| Model | AIC value |
|---|---|
| OLS | 174.232 |
| Poisson | 174.232 |
| Negative binomial | 174.808 |
| ZINB | 158.96 |

TableS3. Results of the GLM evaluating the larval density, resource volume and sex on larval development time of *D. suzukii*. Significant effects ($p<0.05$) are in bold.

| | Sum Sq | df | F | p |
|---|---|---|---|---|
| Density | 77.98 | 4 | 32.437 | **<0.001** |
| Volume | 30.43 | 1 | 50.632 | **<0.001** |
| Block | 49.79 | 1 | 82.843 | **<0.001** |
| Sex | 28.57 | 1 | 47.53 | **<0.001** |
| Volume × Density | 3.36 | 4 | 1.397 | 0.23 |
| Volume × Block | 22.72 | 1 | 37.803 | **<0.001** |
| Density × Block | 25.91 | 4 | 10.778 | **<0.001** |
| Density × Sex | 1.92 | 4 | 0.797 | 0.53 |
| Volume × Sex | 2.29 | 1 | 3.811 | 0.05 |
| Block × Sex | 0.16 | 1 | 0.263 | 0.61 |

TableS4. Comparisons of the resource volume and block type on the larval development time according to the Least Significant Difference post-hoc test. Significant effects ($p<0.05$) are in bold.

| Block | Volume | estimate | z | p |
|---|---|---|---|---|
| 1 | 2-5 | 0.761 | 9.552 | **<0.0001** |
| 2 | 2-5 | 0.11 | 1.334 | 0.18 |

| Volume | Block | estimate | z | p |
|---|---|---|---|---|
| 2 | 1-2 | 0.884 | 10.967 | **<0.0001** |
| 5 | 1-2 | 0.232 | 2.85 | **<0.01** |

TableS5. Comparisons of the density on the larval development time according to the Least Significant Difference post-hoc test. Significant effects (p<0.05) are in bold.

| Block | Density | estimate | z | p |
|---|---|---|---|---|
| 1 | 5-1 | -0.36 | -2.317 | **<0.05** |
| 1 | 10-1 | -1.017 | -6.734 | **<0.0001** |
| 1 | 20-1 | -0.993 | -7.437 | **<0.0001** |
| 1 | 50-1 | -0.944 | -8.252 | **<0.0001** |
| 2 | 5-1 | -0.414 | -2.501 | **<0.05** |
| 2 | 10-1 | -1.002 | -6.49 | **<0.0001** |
| 2 | 20-1 | -0.842 | -6.193 | **<0.0001** |
| 2 | 50-1 | -0.199 | -1.533 | 0.12 |

TableS6. Comparisons of the block type on development time according to the Least Significant Difference post-hoc test. Significant effects (p<0.05) are in bold.

| Density | Block | estimate | z | p |
|---|---|---|---|---|
| 1 | 1-2 | 0.729 | 4.731 | **<0.0001** |
| 5 | 1-2 | 0.784 | 4.664 | **<0.0001** |
| 10 | 1-2 | 0.714 | 4.69 | **<0.0001** |
| 20 | 1-2 | 0.579 | 5.175 | **<0.0001** |
| 50 | 1-2 | -0.016 | -0.199 | 0.84 |

TableS7. Results of the GLMM evaluating the larval density and resource volume on microbial colony counts present in the medium where *D. suzukii* larvae develop. Significant effects (p<0.05) are in bold.

| | $X^2$ | df | p |
|---|---|---|---|
| Density | 19.116 | 4 | **<0.001** |
| Volume | 0.045 | 1 | 0.83 |
| Medium | 25.739 | 1 | **<0.001** |
| Volume × Density | 6.429 | 4 | 0.17 |
| Volume × Medium | 21.323 | 1 | **<0.001** |
| Density × Medium | 112.858 | 4 | **<0.001** |

TableS8. Comparisons of the larval density and growth medium on microbial colony counts according to the Tukey post-hoc test. Significant effects (p<0.05) are in bold.

| Density/Medium | estimate | z | p |
|---|---|---|---|

| | | | |
|---|---|---|---|
| 5.LB – 1.LB | 3.103 | 2.054 | 0.38 |
| 10.LB – 1.LB | 5.965 | 4.023 | **<0.001** |
| 20.LB – 1.LB | 6.968 | 4.7 | **<0.001** |
| 50.LB – 1.LB | 6.478 | 4.262 | **<0.001** |
| 1.TSA – 1.LB | -0.431 | -18.085 | **<0.001** |
| 5.TSA – 1.LB | 2.797 | 1.852 | 0.53 |
| 10.TSA – 1.LB | 6.014 | 4.056 | **<0.001** |
| 20.TSA – 1.LB | 6.822 | 4.602 | **<0.001** |
| 50.TSA – 1.LB | 6.454 | 4.246 | **<0.001** |
| 10.LB – 5.LB | 2.862 | 2.063 | 0.37 |
| 20.LB – 5.LB | 3.865 | 2.792 | 0.06 |
| 50.LB – 5.LB | 3.375 | 2.368 | 0.2 |
| 1.TSA – 5.LB | -3.534 | -2.34 | 0.21 |
| 5.TSA – 5.LB | -0.305 | -27.468 | **<0.001** |
| 10.TSA – 5.LB | 2.911 | 2.098 | 0.35 |
| 20.TSA – 5.LB | 3.719 | 2.687 | 0.09 |
| 50.TSA – 5.LB | 3.351 | 2.351 | 0.21 |
| 20.LB – 10.LB | 1.003 | 0.749 | 0.99 |
| 50.LB – 10.LB | 0.512 | 0.371 | 1 |
| 1.TSA – 10.LB | -6.396 | -4.314 | **<0.001** |
| 5.TSA – 10.LB | -3.168 | -2.284 | 0.24 |
| 10.TSA – 10.LB | 0.048 | 5.39 | **<0.001** |
| 20.TSA – 10.LB | 0.857 | 0.64 | 0.99 |
| 50.TSA – 10.LB | 0.488 | 0.354 | 1 |
| 50.LB – 20.LB | -0.49 | -0.356 | 1 |
| 1.TSA – 20.LB | -7.399 | -4.991 | **<0.001** |
| 5.TSA – 20.LB | -4.171 | -3.013 | **<0.05** |
| 10.TSA – 20.LB | -0.954 | -0.713 | 0.99 |
| 20.TSA – 20.LB | -0.146 | -18.06 | **<0.001** |
| 50.TSA – 20.LB | -0.514 | -0.374 | 1 |
| 1.TSA – 50.LB | -6.909 | -4.545 | **<0.001** |
| 5.TSA – 50.LB | -3.681 | -2.582 | 0.12 |
| 10.TSA – 50.LB | -0.464 | -0.336 | 1 |
| 20.TSA – 50.LB | 0.344 | 0.25 | 1 |
| 50.TSA – 50.LB | -0.024 | -2.531 | 0.13 |
| 5.TSA – 1.TSA | 3.228 | 2.137 | 0.32 |
| 10.TSA – 1.TSA | 6.445 | 4.347 | **<0.001** |
| 20.TSA – 1.TSA | 7.253 | 4.892 | **<0.001** |
| 50.TSA – 1.TSA | 6.885 | 4.529 | **<0.001** |
| 10.TSA – 5.TSA | 3.216 | 2.318 | 0.22 |
| 20.TSA – 5.TSA | 4.025 | 2.908 | **<0.05** |
| 50.TSA – 5.TSA | 3.656 | 2.565 | 0.12 |
| 20.TSA – 10.TSA | 0.808 | 0.604 | 0.99 |
| 50.TSA – 10.TSA | 0.44 | 0.318 | 1 |
| 50.TSA – 20.TSA | -0.368 | -0.268 | 1 |

TableS9. Comparisons of the resource volume and growth medium on microbial colony counts according to the Tukey post-hoc test. Significant effects ($p<0.05$) are in bold.

| **Volume/Medium** | **estimate** | **z** | **p** |
|---|---|---|---|
| 5.LB – 2.LB | 1.258 | 1.206 | 0.54 |

| | | | |
|---|---|---|---|
| 2.TSA – 2.LB | -0.025 | -3.182 | **<0.01** |
| 5.TSA – 2.LB | 1.113 | 1.067 | 0.63 |
| 2.TSA – 5.LB | -1.284 | -1.23 | 0.52 |
| 5.TSA – 5.LB | -0.145 | -26.369 | **<0.001** |
| 5.TSA – 2.TSA | 1.138 | 1.091 | 0.62 |

TableS10. Results of the GLM evaluating the larval density and resource volume on fecundity of *D. suzukii*. Significant effects (p<0.05) are in bold.

| | **Sum Sq** | **df** | **F** | **p** |
|---|---|---|---|---|
| Density | 424.6 | 4 | 4.246 | **<0.01** |
| Volume | 687.2 | 1 | 27.493 | **<0.001** |
| Block | 113.1 | 1 | 4.525 | **<0.05** |
| Volume × Density | 149.8 | 4 | 1.498 | 0.2 |
| Volume × Block | 5.1 | 1 | 0.205 | 0.65 |
| Density × Block | 286.7 | 4 | 2.868 | **<0.05** |

TableS11. Comparisons of the larval density on fecundity according to the Least Significant Difference post-hoc test. Significant effects (p<0.05) are in bold.

| **Block** | **Density** | **estimate** | **z** | **p** |
|---|---|---|---|---|
| 1 | 5-1 | -2.482 | -1.712 | 0.24 |
| 1 | 10-1 | 1.642 | 1.178 | 0.24 |
| 1 | 20-1 | -1.784 | -1.367 | 0.24 |
| 1 | 50-1 | -3.214 | -2.463 | 0.05 |
| 2 | 5-1 | 2.385 | 1.481 | 0.41 |
| 2 | 10-1 | -0.222 | -0.153 | 0.87 |
| 2 | 20-1 | -2.492 | -1.785 | 0.29 |
| 2 | 50-1 | -1.686 | -1.18 | 0.47 |

TableS12. Comparisons of the block type on fecundity according to the Least Significant Difference post-hoc test. Significant effects (p<0.05) are in bold.

| **Density** | **Block** | **estimate** | **z** | **p** |
|---|---|---|---|---|
| 1 | 1-2 | -0.588 | -0.416 | 0.67 |
| 5 | 1-2 | -5.455 | -3.339 | **<0.001** |
| 10 | 1-2 | 1.277 | 0.889 | 0.37 |
| 20 | 1-2 | 0.12 | 0.094 | 0.92 |
| 50 | 1-2 | -2.115 | -1.605 | 0.11 |

TableS13. Results of the GLM evaluating the larval density and resource volume on wings' length of *D. suzukii*. Significant effects (p<0.05) are in bold.

| | **Sum Sq** | **df** | **F** | **p** |
|---|---|---|---|---|
| Density | 540973 | 4 | 31.408 | **<0.001** |
| Volume | 9588 | 1 | 2.226 | 0.13 |
| Block | 1104 | 1 | 0.256 | 0.61 |
| Volume × Density | 81932 | 4 | 4.756 | **<0.01** |
| Volume × Block | 114482 | 1 | 26.586 | **<0.001** |

| | | | | |
|---|---|---|---|---|
| Density × Block | 21897 | 4 | 1.271 | 0.28 |

TableS14. Comparisons of the larval density on wings' length according to the Least Significant Difference post-hoc test. Significant effects ($p<0.05$) are in bold.

| **Volume** | **Density** | **estimate** | **z** | **p** |
|---|---|---|---|---|
| 2 | 5-1 | -9.24 | -0.461 | 0.64 |
| 2 | 10-1 | -34.28 | -1.909 | 0.11 |
| 2 | 20-1 | -88.36 | -5.211 | **<0.0001** |
| 2 | 50-1 | -168.64 | -9.945 | **<0.0001** |
| 5 | 5-1 | 14.23 | 0.706 | 0.64 |
| 5 | 10-1 | 9.17 | 0.471 | 0.64 |
| 5 | 20-1 | -33.88 | -1.83 | 0.2 |
| 5 | 50-1 | -63.67 | -3.36 | **<0.01** |

TableS15. Comparisons of the resource volume on wings' length according to the Least Significant Difference post-hoc test. Significant effects ($p<0.05$) are in bold.

| **Density** | **Volume** | **estimate** | **z** | **p** |
|---|---|---|---|---|
| 1 | 2-5 | 32.8 | 1.761 | 0.07 |
| 5 | 2-5 | 9.33 | 0.435 | 0.66 |
| 10 | 2-5 | -10.65 | -0.565 | 0.57 |
| 20 | 2-5 | -21.69 | -1.29 | 0.19 |
| 50 | 2-5 | -72.17 | -4.17 | **<0.0001** |

TableS16. Comparisons of the resource volume and block type on the wings' length according to the Least Significant Difference post-hoc test. Significant effects ($p<0.05$) are in bold.

| **Block** | **Volume** | **estimate** | **z** | **p** |
|---|---|---|---|---|
| 1 | 2-5 | -55 | -4.824 | **<0.0001** |
| 2 | 2-5 | 30.1 | 2.49 | **<0.05** |

| **Volume** | **Block** | **estimate** | **z** | **p** |
|---|---|---|---|---|
| 2 | 1-2 | -38.3 | -3.32 | **<0.001** |
| 5 | 1-2 | 46.8 | 3.923 | **<0.001** |

TableS17. Results of the GLM evaluating the larval density and resource volume on wings' width of *D. suzukii*. Significant effects ($p<0.05$) are in bold.

| | **Sum Sq** | **df** | **F** | **p** |
|---|---|---|---|---|
| Density | 201876 | 4 | 28.872 | **<0.001** |
| Volume | 6890 | 1 | 3.942 | **<0.05** |
| Block | 1068 | 1 | 0.611 | 0.43 |
| Volume × Density | 35752 | 4 | 5.113 | **<0.001** |
| Volume × Block | 25961 | 1 | 14.851 | **<0.001** |
| Density × Block | 1091 | 4 | 0.156 | 0.96 |

TableS18. Comparisons of the larval density on wings' width according to the Least Significant Difference post-hoc test. Significant effects (p<0.05) are in bold.

| Volume | Density | estimate | z | p |
|---|---|---|---|---|
| 2 | 5-1 | -15.18 | -1.188 | 0.23 |
| 2 | 10-1 | -32.37 | -2.829 | **<0.01** |
| 2 | 20-1 | -61.48 | -5.691 | **<0.0001** |
| 2 | 50-1 | -110.48 | -10.225 | **<0.0001** |
| 5 | 5-1 | -3.34 | -0.26 | 0.79 |
| 5 | 10-1 | -6.63 | -0.534 | 0.79 |
| 5 | 20-1 | -31.06 | -2.633 | **<0.05** |
| 5 | 50-1 | -42.06 | -3.484 | **<0.01** |

TableS19. Comparisons of the resource volume on wings' width according to the Least Significant Difference post-hoc test. Significant effects (p<0.05) are in bold.

| Density | Volume | estimate | z | p |
|---|---|---|---|---|
| 1 | 2-5 | 16.71 | 1.408 | 0.16 |
| 5 | 2-5 | 4.87 | 0.356 | 0.72 |
| 10 | 2-5 | -9.03 | -0.752 | 0.45 |
| 20 | 2-5 | -13.72 | -1.281 | 0.2 |
| 50 | 2-5 | -51.71 | -4.692 | **<0.0001** |

TableS20. Comparisons of the resource volume and block type on the wings' width according to the Least Significant Difference post-hoc test. Significant effects (p<0.05) are in bold.

| Block | Volume | estimate | z | p |
|---|---|---|---|---|
| 1 | 2-5 | -30.84 | -4.243 | **<0.0001** |
| 2 | 2-5 | 9.69 | 1.258 | 0.21 |

| Volume | Block | estimate | z | p |
|---|---|---|---|---|
| 2 | 1-2 | -24.4 | -3.32 | **<0.001** |
| 5 | 1-2 | 16.1 | 2.12 | **<0.05** |

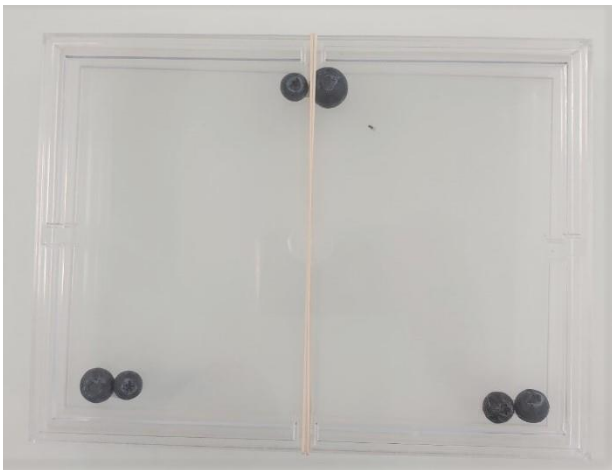

FigureS1. Experimental set up for the oviposition assays on blueberries. Three groups of 2 blueberries were placed at equal distance from each other in a box (23.8cm x 17.8cm x 2cm).

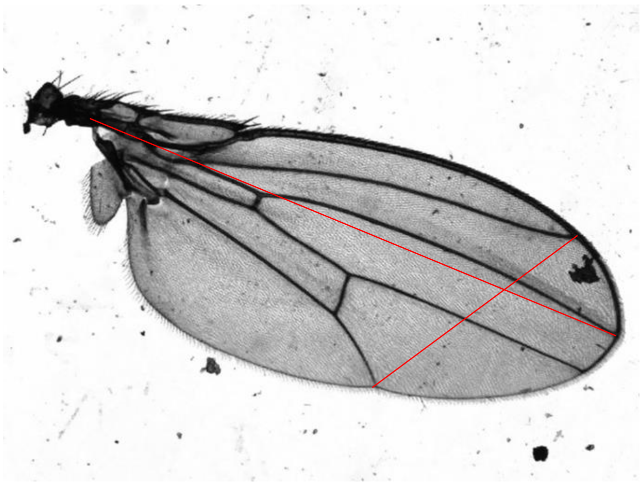

FigureS2. Measures done for the wing size. The two red lines indicated the measures done.

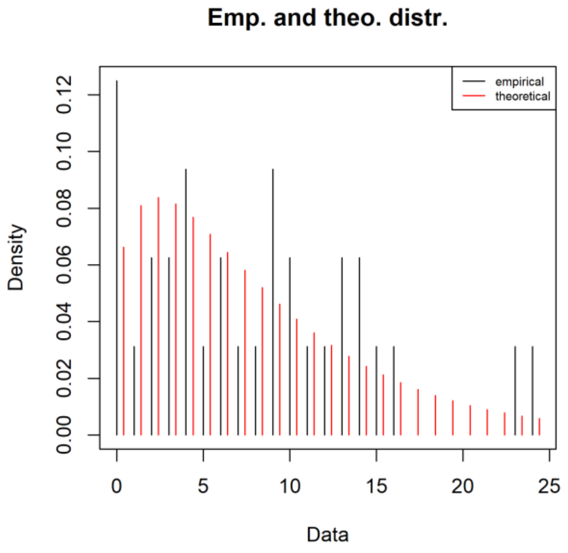


FigureS3. Comparison between the empirical and theoretical (zero-inflated negative binomial regression, ZINB) distribution of *D. suzukii* eggs deposited per infested fruit.

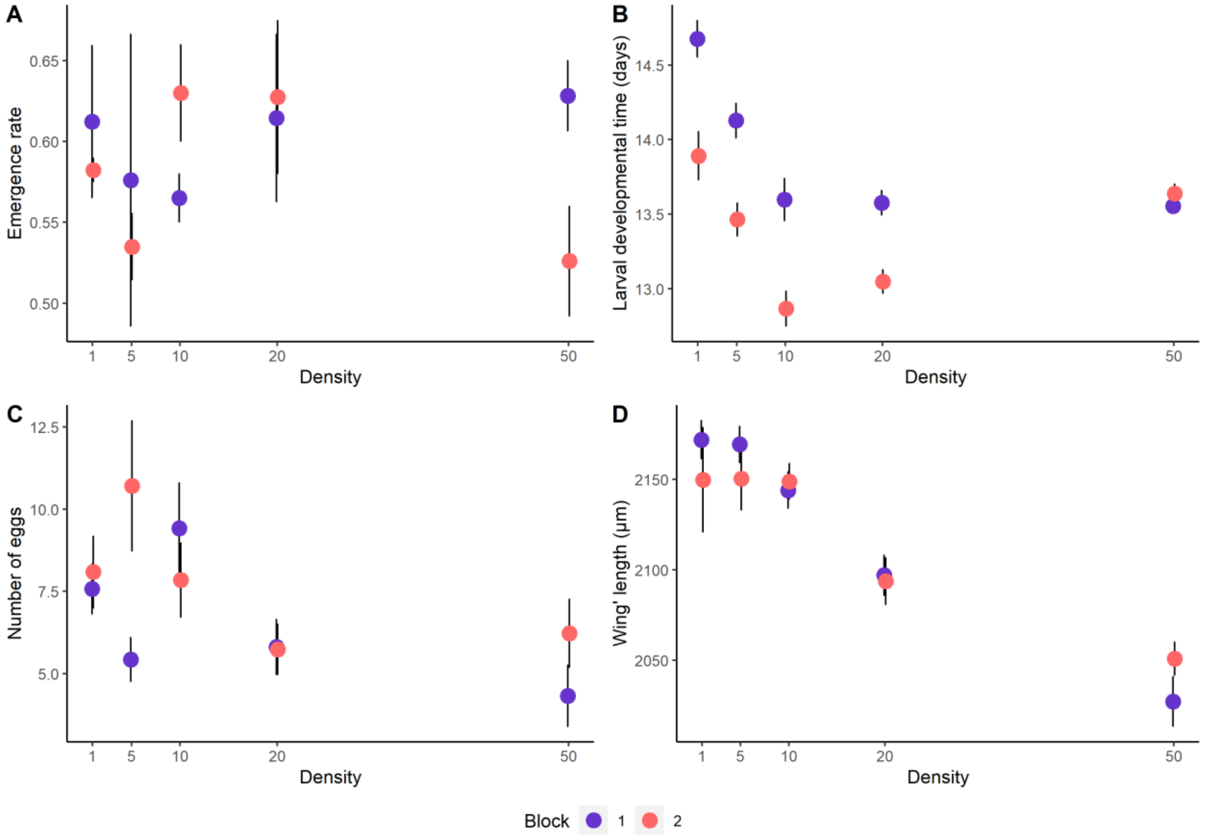


FigureS4. Effect of larval density and block (1 and 2) on immediate (A, B) and delayed (C, D) life-history traits (mean ± SE) of *D. suzukii*. Panel (A) shows emergence rate. Panel (B) shows the developmental time for larvae to pupation in days. Panel (C) shows fecundity measured as the number of eggs. Panel (D) shows wings' length.

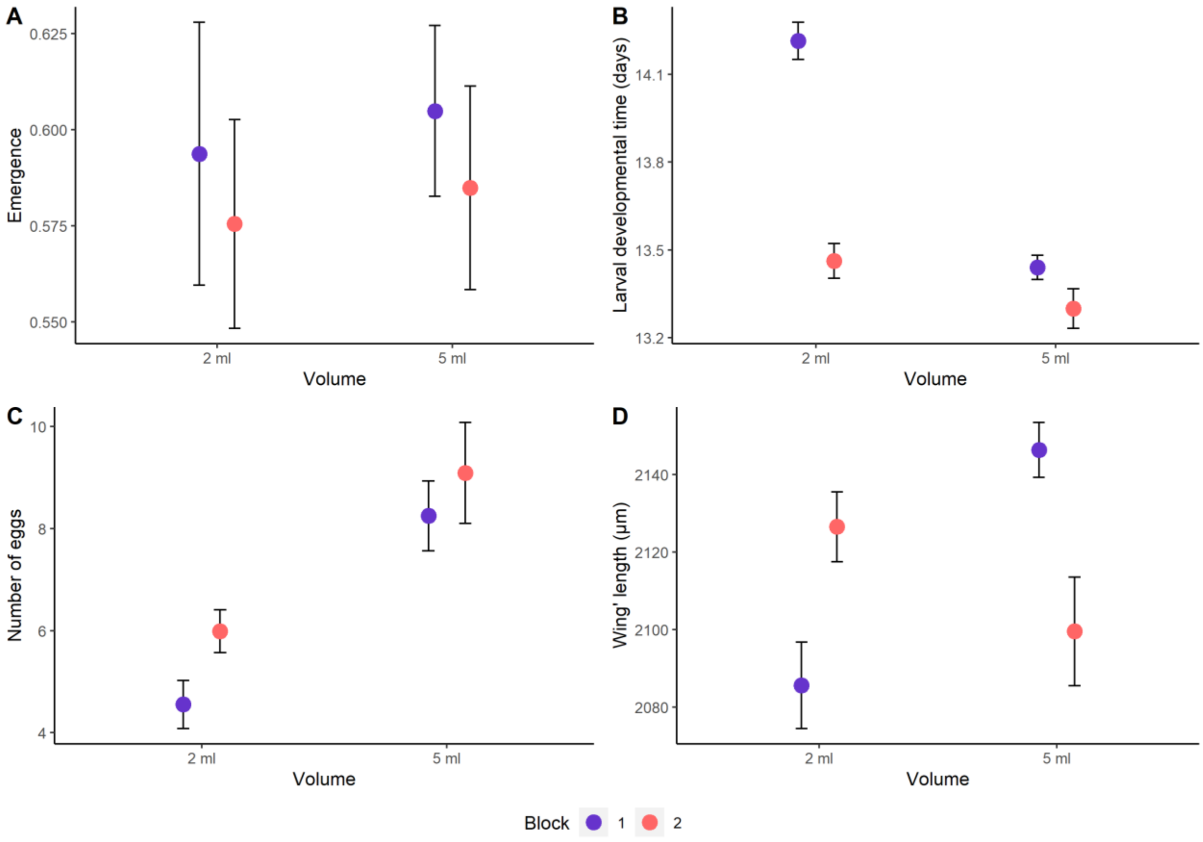


FigureS5. Effect of resource volume (2 and 5 mL of food medium) and block (1 and 2) on immediate (A, B) and delayed (C, D) life-history traits (mean ± SE) of *D. suzukii*. Panel (A) shows emergence rate. Panel (B) shows the developmental time for larvae to pupation in days. Panel (C) shows fecundity measured as the number of eggs. Panel (D) shows wings' length.

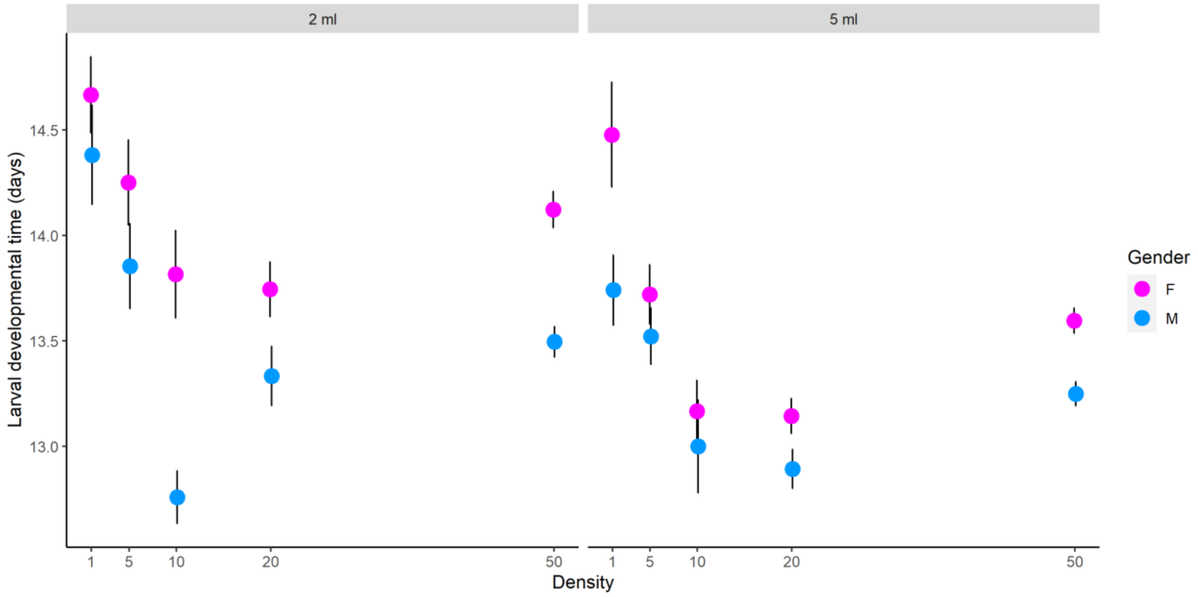


FigureS6. Effect of larval density and resource volume (2 and 5 mL of food medium) on larval developmental time (mean ± SE) between females and males of *D. suzukii*.

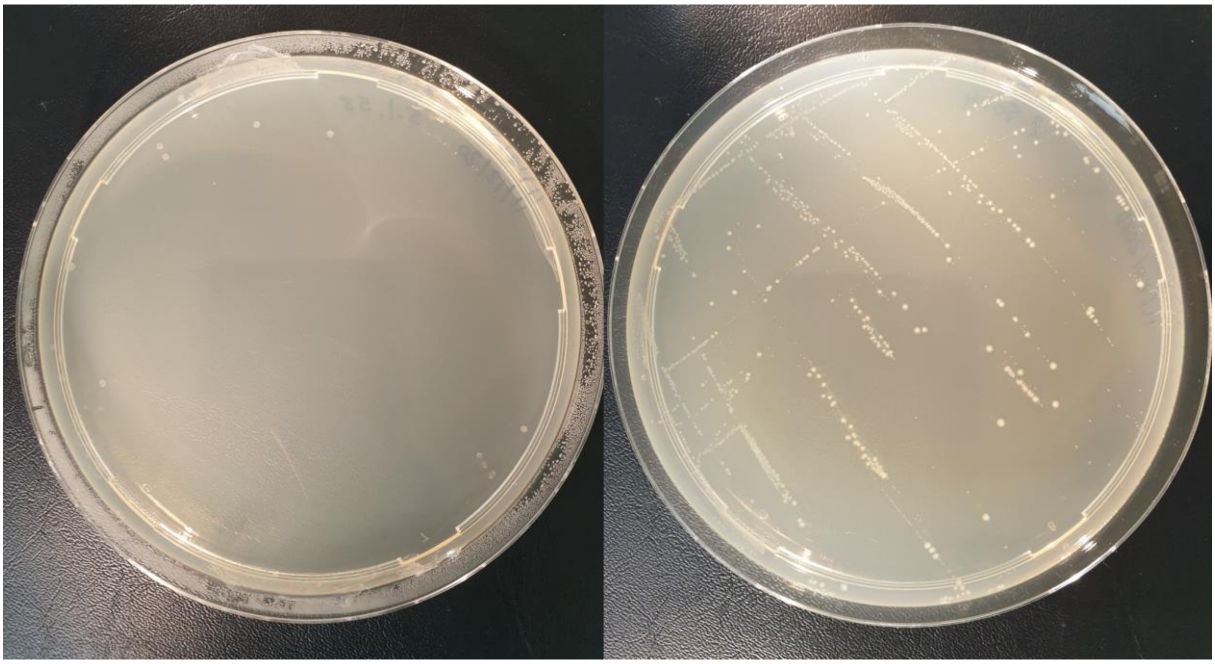

FigureS7. Samples of microbial colonies inoculated in LB medium. On the left, a plate with a few microbial growth and on the right, a medium with a greater number of microbial colonies are shown.

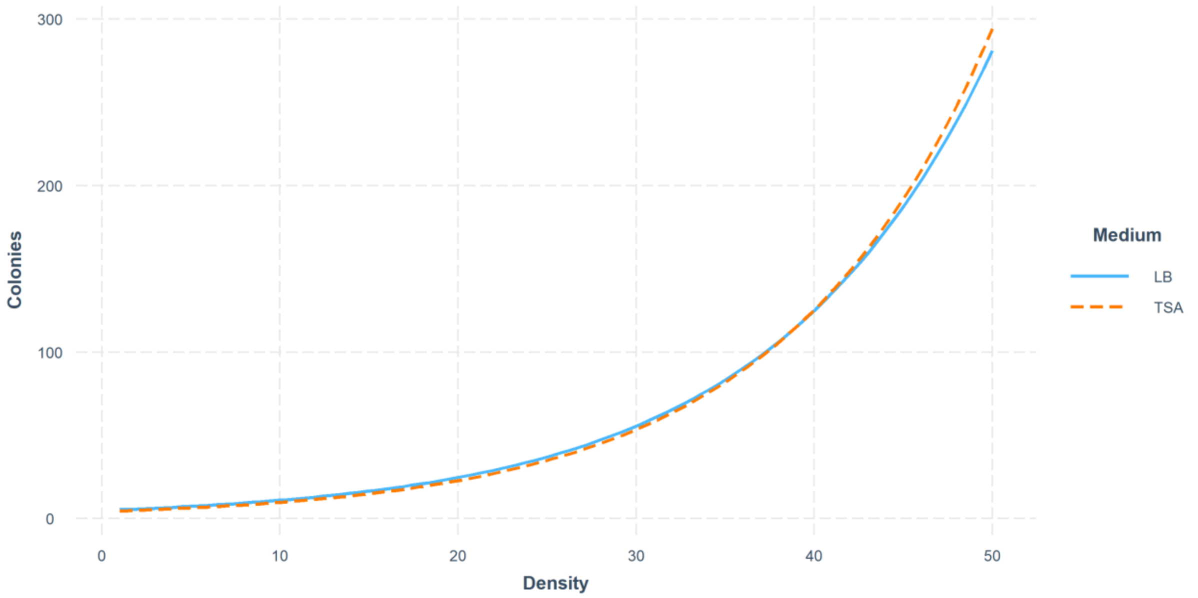


FigureS8. Effect of the interaction between larval density and growth medium on the number of microbial colonies.

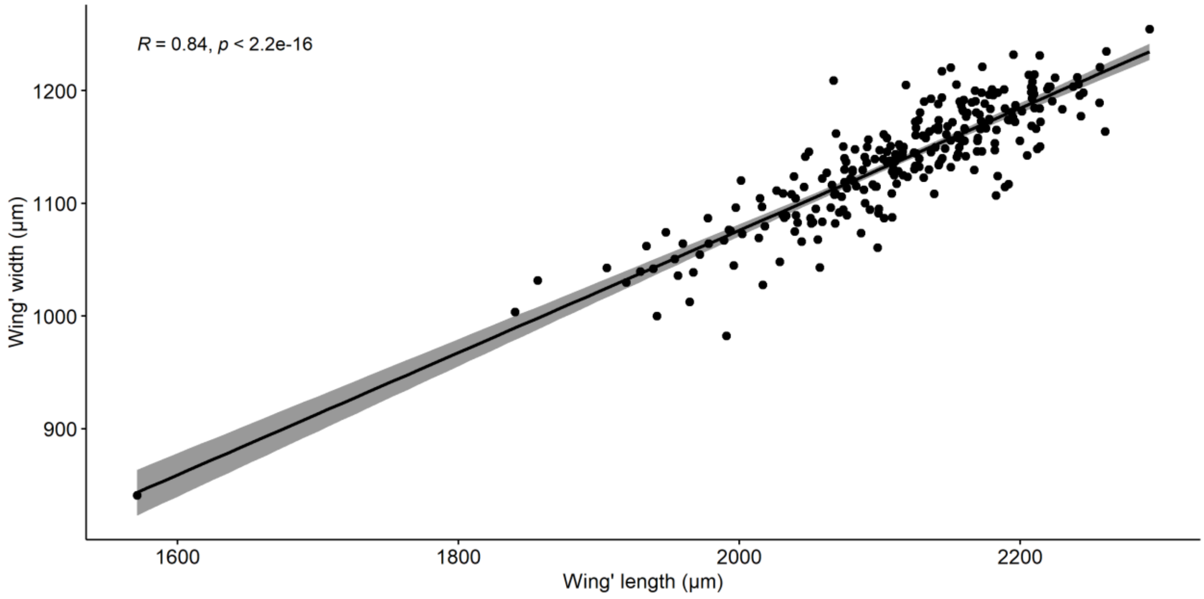


FigureS9. Spearman' correlation coefficient (R) and linear regression line between wings' length and wings' width means. Each point corresponds to one individual (n=257).

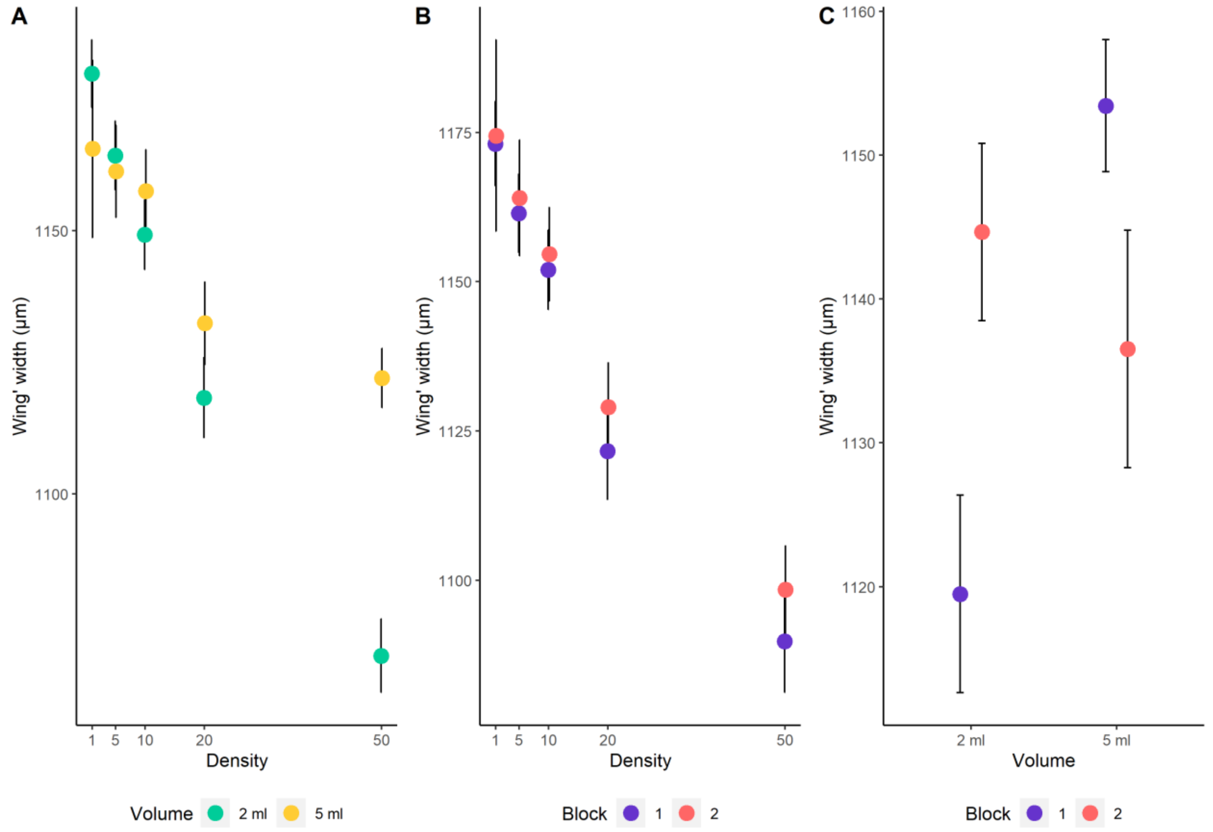


FigureS10. Effect of larval density, resource volume and block on wings' width (mean ± SE) of *D. suzukii*. Panel (A) shows the interaction between larval density and resource volume. Panel (B) shows the interaction between density and block. And panel (C) shows the interaction between volume and block.
